# The Gut Microbiome of the Eastern Spruce Budworm Does Not Influence Larval Growth or Survival

**DOI:** 10.1101/330928

**Authors:** Melbert T. Schwarz, Daniel Kneeshaw, Steven W. Kembel

## Abstract

Microbial communities have been shown to play an important role for host health in mammals, especially humans. It is thought that microbes could play an equally important role in other animal hosts such as insects. A growing body of evidence seems to support this, however most of the research effort in understanding host-microbe interactions in insects has been focused on a few well-studied groups such as bees, cockroaches and termites. We studied the effects of the gut-associated microbial community on the growth and survival of the eastern spruce budworm *Choristoneura fumiferana*, an economically important lepidopteran forest pest in eastern Canada and the northeastern United States. Contrary to our expectations, the gut microbial community of spruce budworm larvae does not appear to influence host growth or survival. Our results agree with the hypothesis that lepidopteran larvae lack resident microbial communities and are not nutritionally dependent on bacterial symbionts.

## INTRODUCTION

The eastern spruce budworm (*Choristoneura fumiferana*) is a forest pest native to the north eastern United States and eastern Canada that undergoes epidemic population outbreaks every 30-40 years. During these population outbreaks, lasting approximately ten years, millions of hectares of balsam fir (*Abies balsamea*) and spruce (*Picea spp*.) trees are defoliated (Boulanger & Arseneault, 2004; Burton, Svoboda, Kneeshaw, & Gottschalk, 2015; Royama et al., 2005; Sainte-Marie, Kneeshaw, MacLean, & Hennigar, 2015). Consequently spruce budworm has significant effects on forest productivity (MacLean, 1984) and is an economically important defoliator in coniferous forests (Fournier, Bauce, Dupontb, & Berthiaumea, 2010). While the role of factors such as landscape context, parasitoids and other predators, and forest management practices have been hypothesized to play a role in governing spruce budworm health and population dynamics (Régnière & Nealis, 2007; Robert, Kneeshaw, & Sturtevant, 2012), much less is known about the importance of the microbial life associated with spruce budworm for their growth, population dynamics, and outbreak status.

Insects are associated with diverse communities of microorganisms including bacteria and fungi. The collective set of microbial genomes associated with a host, the microbiome, has a much greater functional diversity than the eukaryotic host genome (Lapierre & Gogarten, 2009). One important function of the insect gut microbiome is its potential to aid in digestion by breaking down compounds the host cannot digest (Douglas, 2009; Feldhaar, 2011). Thus, the microbiome can act as an extension of the host gut. This is particularly important when considering the spruce budworm because it feeds on conifer needles that are difficult to digest and contain defensive compounds such as terpenes (Mumm & Hilker, 2006), such that the microbiome could potentially aid the host by suppressing plant defenses or helping the host detoxify defensive secondary chemicals.

Drawing generalizations about the extent to which insect microbiota are important to their hosts is difficult due in large part to the morphological, physiological, and behavioral variation among insects. Different physiology, life histories, and feeding strategies can all influence how microbes interact with their host (Philipp Engel & Moran, 2013). There is evidence to support the assumption that gut microbial symbiosis is important in a number of insects (De Souza et al., 2013; Emery, Schmidt, & Engel, 2017; P. Engel, Martinson, & Moran, 2012; Philipp Engel & Moran, 2013; Koch & Schmid-Hempel, 2011; Kwong, Mancenido, & Moran, 2017; Prado, Hung, Daugherty, & Almeida, 2010; Rosengaus, Zecher, Schultheis, Brucker, & Bordenstein, 2011). For example, termites depend on specialized bacteria or flagellates to allow them to digest cellulose (Douglas, 2009; Philipp Engel & Moran, 2013; Rosengaus et al., 2011), and various species of bees benefit greatly from their associated microbiota (Philipp Engel & Moran, 2013; De Souza et al., 2013; Rosengaus et al., 2011; Emery, Schmidt, & Engel, 2017). Bee-associated microbes have been shown to contribute to immune function as well as to aid in nutrition through mediating the digestion of pectin (P. Engel et al., 2012; Kwong et al., 2017).

The lepidopteran microbiome has been described as a very simple microbial community compared to other insects (Belda et al., 2011; Brinkmann, Martens, & Tebbe, 2008; Broderick, Raffa, Goodman, & Handelsman, 2004; Landry, Comeau, Derome, Cusson, & Levesque, 2015; Mason & Raffa, 2014; Robinson, Schloss, Ramos, Raffa, & Handelsman, 2010; Tang et al., 2012; Xiang et al., 2006). One reason that lepidopteran gut microbial communities tend to be simpler than other insects is that the lepidopteran larval gut is a simple tube without any specialized structure for microbial cultivation as is seen in termite guts (Philipp Engel & Moran, 2013). Another unique aspect of the lepidopteran larval gut is that unlike most insect midguts which are acidic and range in pH between 4-7, lepidopteran midguts are highly alkaline ranging from pH 8-12 (Belda et al., 2011; Broderick et al., 2004; Philipp Engel & Moran, 2013; Tang et al., 2012). The alkaline nature of the spruce budworm gut could provide some advantage in digesting acidic conifer needles.

The objectives of this study were to determine if the gut microbiota associated with spruce budworm larvae influence larval growth rates and survival., and to determine if the eastern spruce budworm has a gut microbiome that is distinct from the microbial assemblages associated with its diet. We also sought to quantify the effects that antibiotics have on spruce budworm gut microbial diversity and community composition to better understand the role of the gut microbiota in regulating spruce budworm growth and survival.

Given the challenges associated with a diet of conifer needles, we hypothesized that the eastern spruce budworm has a resident gut microbiome that contributes to larval growth and survival. Thus, we further hypothesized that the disturbance of microbial communities with antibiotics will negatively influence spruce budworm larval growth and survival. We also hypothesized that the use of antibiotics will both reduce diversity of the microbial communities associated with diet, guts, and frass, and would significantly alter the composition of the spruce budworm gut microbial community in a way that would negatively influence larval growth and survival.

## MATERIALS AND METHODS

### Insect rearing

We acquired approximately 1,000 spruce budworm second instar larvae that had completed diapause from the Insect Production Services at the Great Lakes Forestry Centre, (Sault Ste. Marie, ON, Canada). Larvae were packaged between a sheet of parafilm and a sheet of cheese cloth and stored at 4°C prior to the start of the experiment. Sections of the parafilm containing approximately 30-40 larvae were cut using scissors sterilized for 5 seconds with 70% ethanol and placed on cups of synthetic McMoran diet (McMorran, 1965) containing antibiotics purchased from the Insect Production Services (Sault Ste. Marie, ON, Canada) in previously autoclaved magenta boxes. Larvae were allowed to emerge from their hibernacula and feed on the common diet for one week. The purpose of rearing larvae on a common diet for the first week was twofold: to ensure larvae were large enough to successfully eat foliage, and so that all larvae started to feed on the same food to control for variation in the starting microbiota among second instar larvae. Throughout the experiment larvae were maintained at 24°C at 60% relative humidity under a 16h:8h light:dark cycle.

After 1 week of feeding on the common diet, 200 larvae were randomly selected and split equally among 5 treatments (n=40): artificial diet with antibiotics, black spruce (*Picea mariana*) foliage treated with antibiotics, untreated spruce foliage, balsam fir (*Abies balsamea*) foliage treated with antibiotics, and untreated balsam fir foliage. Each replicate consisted of an individual larva in an autoclaved magenta box. Spruce foliage was collected from saplings housed in the greenhouse at the Université du Québec à Montréal and stored at −20°C for approximately 4 weeks. Fir foliage was collected from trees near Baie Comeau, Québec and stored in sterile bags at −20°C for 4 weeks. In both cases we took care to use only foliage that had fresh growth. Foliage was placed in a 2ml microcentrifuge tube filled with sterile water, or 50 μg/ml streptomycin for antibiotic treatments, and sealed with parafilm to reduce desiccation of the cut foliage during the experiment. For antibiotic treatments, a 1500 ppm solution of methyl paraben and a 50 μg/ml solution of streptomycin were each sprayed on the foliage every other day. Untreated foliage was not manipulated other than placing the cut stem in a microcentrifuge tube containing sterile water.

### Health assessment: measuring larval growth and survival

Larval health was assessed every other day by measuring larval weight and calculating growth rates. Overall survival was calculated as well. We chose larval weight as a measure of health because it is often used as a measure of fitness in pupae and therefore can also be used as a representation of overall health (Hammer, Janzen, Hallwachs, Jaffe, & Fierer, 2017). We removed each larva from its magenta box using a fine paintbrush, placed it on a sterile weigh boat, and recorded the mass. Re-application of antibiotics on foliage occurred at this time via spray bottle. All work was done in an ethanol sterilized fume hood. The paintbrush used to manipulate the larvae was sterilized for 5 seconds with 70% ethanol between each replicate. Larvae that were dead at the time of weighing were discarded.

### Sample collection

We collected sixth-instar larvae just prior to pupation, placed them in microcentrifuge tubes, and left them at room temperature for 4 hours before freezing at −80°C. This was to allow for any remaining food to pass through their guts, providing us with a more accurate approximation of the true gut microbiota as opposed to microbes that simply pass through the gut along with the food. Larval midguts were extracted from surviving individuals, using forceps and scissors sterilized with 70% ethanol, by cutting the posterior and anterior ends of the individual off to separate the midgut from the hindgut and foregut; remaining midgut was extracted from the larva using the forceps. Extracted guts were placed directly in MoBio PowerSoil bead beating tubes (Qiagen) and stored at −20°C until nucleic acid extraction. We chose to sample the midgut because in many insects the majority of digestion and absorption happens in the midgut (Philipp Engel & Moran, 2013). Additionally, the relatively fast gut passage through the larval budworm, along with a lack of specialized gut structure, would indicate that the hindgut microbial community is not critical in digestion via microbial fermentation for these insects. Finally, the midgut of spruce budworm has been sampled in a previous study investigating rearing effects on the spruce budworm microbiome (Landry et al., 2015).

We sampled frass and foliage samples twice during the experiment, once 7 days after exposure to treatments and again after 14 days when larvae were also collected. Foliage was collected by taking 5 needles with ethanol sterilized forceps, placed in microcentrifuge tubes, and immediately frozen at −80°C. Frass was collected from the bottom of the magenta box, placed in microcentrifuge tubes, and immediately frozen at −80°C. Samples were then assessed for microbial community diversity and composition following DNA sequencing.

We sampled frass and foliage communities along with the gut microbiota. Using these three communities to determine how the relative abundance of microbes changes from the source (foliage) through an environmental filter (gut) and by comparing gut communities with frass communities (or foliage communities) makes it possible to determine which taxa are able to persist in the gut versus which taxa simply pass through the larval gut.

### DNA extraction and processing

We extracted DNA from the midguts of all surviving larvae (n=96). In addition, 10 individuals were selected randomly from each of the 5 treatment groups to extract DNA from foliage or synthetic diet (n=101) and frass (n=99), both collected at each of the two time points. All genomic DNA from the guts, foliage, and frass was extracted using the MoBio PowerSoil DNA extraction kit (Qiagen). We used a slightly altered protocol, as described below, in order to increase DNA yields. Guts were homogenized by vortexing for 10 minutes in the provided PowerSoil bead beating tubes and centrifuged at room temperature for 1 min at 10,000g. The supernatant was transferred to a sterile 2 ml microcentrifuge tube and sonicated with the Bioruptor UCD-200 sonicator (Diagenode) for 1 min on the low setting (160W at 20kHz) for 5 min. After sonication the DNA extraction proceeded as per the manufacturer’s instructions.

Foliage and synthetic diet samples were placed in thick walled 2 ml tubes with three 2.3mm diameter stainless steel beads (BioSpec Products, Bartlesville, OK, USA) and 250 μl of the PowerSoil bead tube buffer. Diets were homogenized using a MiniBead Beadbeater-16 (BioSpec Products, Bartlesville.) for 1.5 minutes. The remaining buffer from the bead beating tube was added to the resulting homogenate, sonicated at the high setting (320W at 20kHz) for 2 minutes and re-introduced to the bead beating tubes. Frass samples were sonicated for 2 minutes at the high setting (320W at 20kHz) for 2 minutes with 250 μl of the bead beating buffer. Following sonication, samples were transferred back to the bead beating tubes. For diet and frass samples, after sonication the DNA extraction was performed following the manufacturer’s instructions.

Following DNA extractions all samples were cleaned using the Zymo OneStep-96 PCR inhibitor removal kit. Polymerase chain reaction (PCR) was used to amplify the 16S rRNA gene from the extracted DNA. We used the chloroplast excluding primers (799F and 1115R) (Chelius & Triplett, 2001) to target the V5-V6 region of the 16S rRNA gene. Each primer also contained 1 of 20 unique bar codes and an Illumina adaptor to allow sequences to bind to the flow cell of the MiSeq sequencer. PCR was performed using 25 μl reactions prepared with 1 μl genomic DNA diluted 1:10 in molecular-grade water, 5 μl 5x HF buffer (Thermo Scientific), 0.5 μl dNTP’s (10 μM each), 0.5 μl forward and reverse primer (10 μM each), 0.75 μl DMSO, 0.25μl Phusion

HotStart II polymerase (Thermo Scientific), and 16.5 μl molecular-grade water. Each reaction began with 30 seconds of denaturation at 98°C followed by 35 cycles of: 15s at 98°C, 30s at 64°C, 30s at 72°C, and a final elongation step at 72°C for 10 minutes. Each PCR included a positive control and a negative control that were verified using gel electrophoresis on an agarose gel prior to sequencing. Amplicons were cleaned and normalized to 0.55 ng/μl using the Invitrogen SequelPrep normalization plate kit. After normalization equal volumes of amplicon DNA per sample were pooled and sequenced. In addition to sequencing experimental samples, positive and negative controls from each PCR were sequenced.

### Amplicon sequencing

We sequenced 16S rRNA gene amplicons using the Illumina MiSeq platform using V3 chemistry. After sequencing, we first trimmed Illumina adapters from our sequences using the program BBduk version 35.76 (https://sourceforge.net/projects/bbmap) and created paired end sequences using PEAR version 0.9.5 (Zhang, Kobert, Flouri, & Stamatakis, 2014). The resulting paired end sequences were analyzed using DADA2 v 1.9.3 (Benjamin J. Callahan et al., 2016). Sequences were trimmed by truncating forward and reverse reads at 250 and 200 base pairs respectively and removing sequences with more than 2 expected errors or any ambiguous bases. Following trimming, error rates were estimated for both forward and reverse reads using the DADA2 error estimation model. Using the calculated error rates, DADA2 was then used to remove sequencing errors from the dataset, merge paired ends, and infer exact sequence variants (ESV). Finally, chimeric sequences were removed. Except where otherwise noted, default settings were used for all bioinformatics analyses. Following quality control steps and the removal of chimeric sequences a final dataset containing 2,955,612 sequences and 1,298 inferred ESVs.

Taxonomy was assigned to each ESV using the DADA2 implementation of the RDP Naïve Bayesian classifier (Ben J. Callahan, Sankaran, Fukuyama, McMurdie, & Holmes, 2016; Wang, Garrity, Tiedje, & Cole, 2007) with the SILVA128 taxonomic database (Quast et al., 2013). In addition an alignment of each ESV’s representative sequence was used to create a phylogeny using the FastTree 2 software (Price, Dehal, & Arkin, 2010). Positive controls were identified as *E. coli* and were distinct from experimental samples as expected. Negative controls had few to zero sequences and did not pass quality control steps. Following quality control, we analyzed community composition, structure, and the effects of gut communities on spruce budworm growth and survival using the statistical software R (R Core Team, 2017).

### Growth and survival analysis

The effects of antibiotic treatment and diet (spruce or fir) on spruce budworm growth and survival were tested using two separate models. A mixed-effects model implemented with the R package nlme (Pinheiro, Bates, DebRoy, Sarkar, & R Core Team, 2017) was used to test for differences in larval growth. Larval weights were log transformed and used as the response variable in the model. Time, antibiotic treatment, diet, and their interactions were used as fixed effects, where time as a fixed effect is an estimate of growth rate, and time nested within individual larvae was used as the random effect for the model. Differences in larval survival were tested using a separate logistic regression with survival as a binary response variable and antibiotic treatment, diet, and their interaction as main effects.

### Community analysis

We tested for differences in community structure, i.e community composition and the relative abundances of the taxa present, as well as diversity both among and within sample types (foliage, guts, frass). When we made comparisons among sample types, data were analyzed as a single dataset so we could ensure that each sample type had equal sampling depth for comparisons. When comparing treatments within sample types we analyzed separate datasets for each sample type that were rarefied separately. Sample types were rarefied separately to ensure the maximum number of sequences could be used in our analysis, allowing for more statistical power when testing within sample type differences.

Prior to our analysis we removed extremely rare ESVs (< 10 sequences) and samples that had fewer than 500 total sequences. A total of 1,020 ESVs remained after removing rare ESVs. When all sample types were analyzed together samples were rarefied to 1,000 sequences. When analyzed separately, gut samples were rarefied to 2,500 sequences per sample while diet and frass samples were rarefied to 1,000 sequences each. We calculated Shannon diversity based on relative abundances of rarefied samples for each data set as a measure of diversity (Haegeman et al., 2013).

Community structure was explored using non-metric multidimensional scaling (NMDS) using weighted UniFrac distances (Lozupone & Knight, 2005). Permutational multivariate ANOVA (PERMANOVA) with 10,000 permutations was used to test for differences in community structure among diets and between antibiotic treatments using weighted UniFrac distance measures. PERMANOVA was implemented using the R package Vegan (Oksanen et al., 2017).

Additionally, the fastest and slowest growing larvae, defined as the larvae in the upper and lower quartile of growth rates respectively, were selected for each experimental group and we compared their gut community structure using PERMANOVA and NMDS.

### Nucleotide sequence accession numbers

16S rRNA amplicon sequence reads, including positive and negative controls, were submitted to the NCBI sequence read archive under the SRA accession number SRP139053 which covers all samples collected for this study between the accession numbers SRX3908565 and SRX3908861 (https://www.ncbi.nlm.nih.gov/sra/SRP139053).

## RESULTS

### Neither diet nor antibiotic treatment affected spruce budworm larval survival

None of our experimental treatments (synthetic diet with antibiotics, fir foliage – the primary host species of the SBW – with and without antibiotics; and black spruce foliage – a secondary host – with and without antibiotics) had a significant effect on larval survival. Diet type did not affect eastern spruce budworm larval survival rates (logistic regression; z= −0.897, p=0.3695), however antibiotic treatment tended to favor survival (logistic regression; z= −1.810, p=0.0702) (Fig S1). The trend of antibiotic treatment favoring survival seems to be driven by the difference between larvae feeding on spruce (30% survival) that was not treated with antibiotics and larvae that fed on synthetic diet (60% survival) that contained antibiotics. Because the synthetic diet (McMorran, 1965) was designed to be optimal for spruce budworm growth and survival, a second logistic regression was performed only on larvae that fed on foliage (with and without antibiotics) and we found that there was no longer a trend of antibiotic treatment on larval survival (logistic regression; z=0.110, p=0.913). Larvae feeding on spruce and fir treated with antibiotics both had 50% survival and larvae feeding on fir without antibiotics had 52.5% survival.

### Diet affects growth rate in spruce budworm larvae

Larvae that fed on synthetic diet grew significantly more (0.14 ± 0.007 g/day) than all other treatment groups (ANOVA; F=61.39, p <0.001) with the next highest growth rate observed in larvae that fed on antibiotic spruce (0.07 ± 0.002 g/day). Because the growth rate of larvae feeding on synthetic diet was, on average, about twice that of all other larvae, we excluded larvae fed on synthetic diet from further analyses of growth.

Antibiotic treatment and time significantly affected the weight of spruce budworm larvae (Table 1, Table S1). In the model, the estimates of time as a main effect represent larval growth rate. The overall differences in growth rate observed between spruce budworm larvae in different experimental groups were due to the antibiotic treatment, however this result must be interpreted carefully because it appears to be largely driven by an interaction between diet and antibiotic treatment rather than a consistent effect of antibiotics on growth rate.

**Table 1.** Growth rates of spruce budworm larvae feeding on spruce and fir needles with and without antibiotics calculated as the estimate of time as fixed effect of a mixed effect model comparing larval weights with time, antibiotic treatment, and diet and their interactions as fixed factors and time nested within individual as random factors. In our model time as a fixed effect represents the growth rate of larvae

| Diet | Treatment | Growth rate (g/day) | Standard error | Lower 95% confidence limit | Upper 95% confidence limit |
| --- | --- | --- | --- | --- | --- |
| Fir | AB | 0.044 | 0.0039 | 0.036 | 0.052 |
| Fir | None | 0.045 | 0.0038 | 0.038 | 0.053 |
| Spruce | AB | 0.073 | 0.0038 | 0.065 | 0.081 |
| Spruce | None | 0.062 | 0.0049 | 0.053 | 0.072 |

Individuals feeding on fir treated with antibiotics grew less than those feeding on antibiotic treated spruce foliage (−0.020 ± 0.005 (mean change ± SE); p<0.0001), and larvae feeding on untreated fir foliage grew less than those feeding on untreated spruce (−0.017 ±.006; p=0.032, Fig. 1, Table 1, Table S2). Growth rates of larvae feeding on antibiotic treated and untreated foliage of the same type (i.e fir or spruce) did not differ. However, there was a significant interaction term in our model between time and diet and a nearly significant interaction term between treatment and diet (Table S2) which suggest that antibiotic treatment in the spruce-fed cohort had a slightly negative effect on growth rate in the first five days of the experiment but a positive effect for the remainder of the experiment compared to the untreated group. Therefore, we argue that differences in growth rate observed among different groups of larvae are due to differences in diet more so than any disturbance in the microbial community caused by the antibiotic treatment.

**Figure 1.**
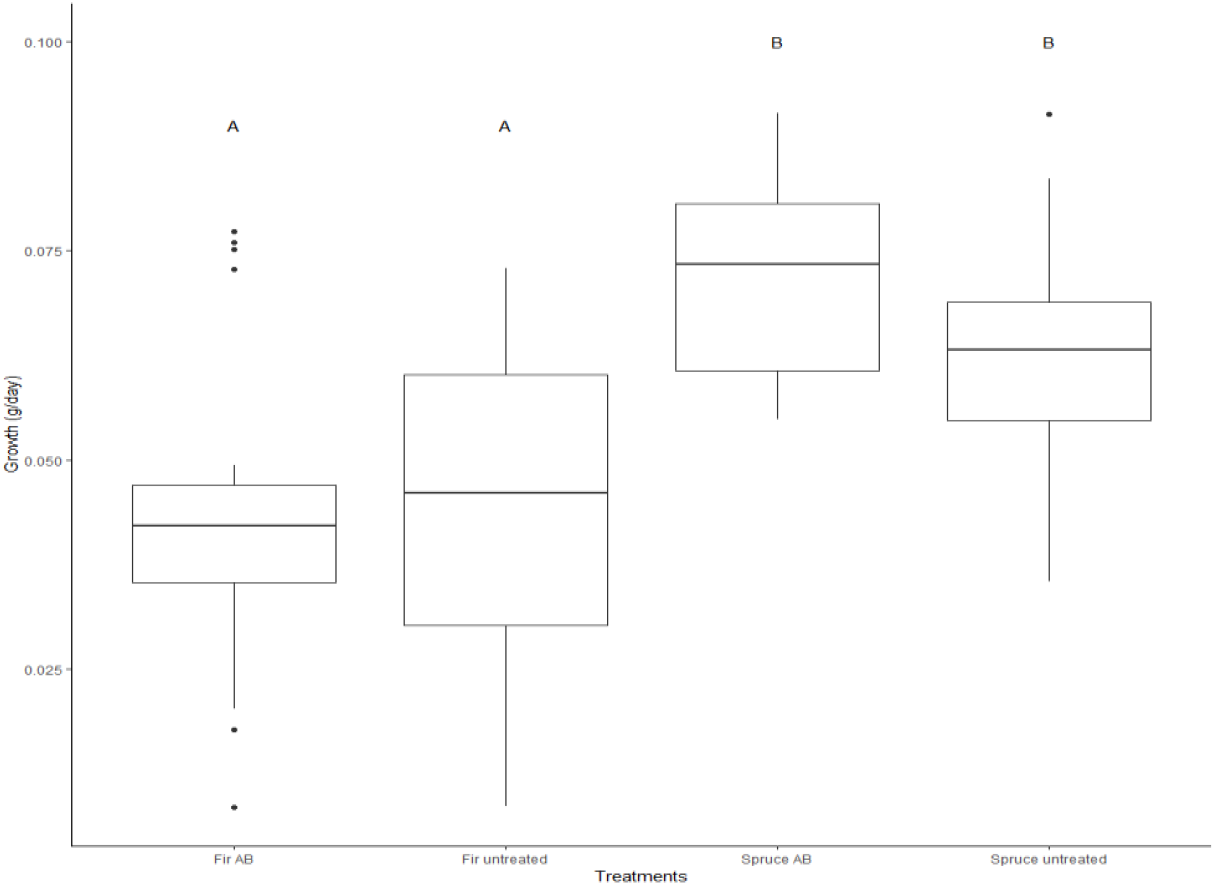
Growth rate (grams/day, ± S.E) of spruce budworm larvae among different diets (spruce versus fir foliage) and antibiotic treatments (AB = antibiotic treated). Letters indicate treatment combinations that differed significantly (p < 0.05) according to a Tukey’s Honest Significant Difference post-hoc test, based on a mixed model (see Methods section for details).

### Gut microbiomes of fast and slow growing larvae did not differ

As an additional way to determine if gut community structure impacts spruce budworm larval growth we compared the gut-associated communities of the fastest and slowest growing larvae (upper and lower quartile of growth rates respectively) in each treatment group. Gut communities of larvae did not differ between fast and slow growers regardless of diet, antibiotic treatment, or distance measure used (PERMANOVA on weighted UniFrac; antibiotic treated fir F=1.89, R^2^=0.32, p=0.10; untreated fir F=0.74, R^2^=0.15, p=0.86; antibiotic spruce F=1.07, R^2^=0.26, p=0.2; untreated spruce F=2.17, R^2^=0.35, p=0.10) further suggesting that the gut microbiome is unimportant for larval growth.

### Antibiotic treatment altered the structure of diet-associated communities but not in larval guts

Here we examined the impact that the antibiotic treatment had on the gut microbiome of spruce budworm larvae independent on the diet type (spruce or fir foliage, or artificial diet). Although we excluded the artificial diet control group from our growth analyses due to the difference between the control group and all of the foliage fed larvae, we included sequencing data from the synthetic diet in our community analyses. This was done to verify whether the microbial community between the common starting diet, which included antibiotics (see methods), and the experimental treatments (spruce or fir foliage treated with and without an antibiotic cocktail) changed over the course of the experiment.

Antibiotic treatment significantly affected the structure (PERMANOVA on weighted UniFrac; F= 3.03, R^2^=0.038, p=0.019) of the larval diet. In this case, structure refers to the composition of the microbial community including the relative abundance of its members, such that it represents both species richness and species evenness. Despite altering the structure of the diet-associated community, antibiotic treatment did not affect the microbial diversity associated with diets (Fig. 2). Because the antibiotic treatment impacted the structure of diet communities but not the diversity, our results show that the antibiotic treatment impacted some bacteria but did not eliminate or select for any specific taxa.

**Figure 2.**
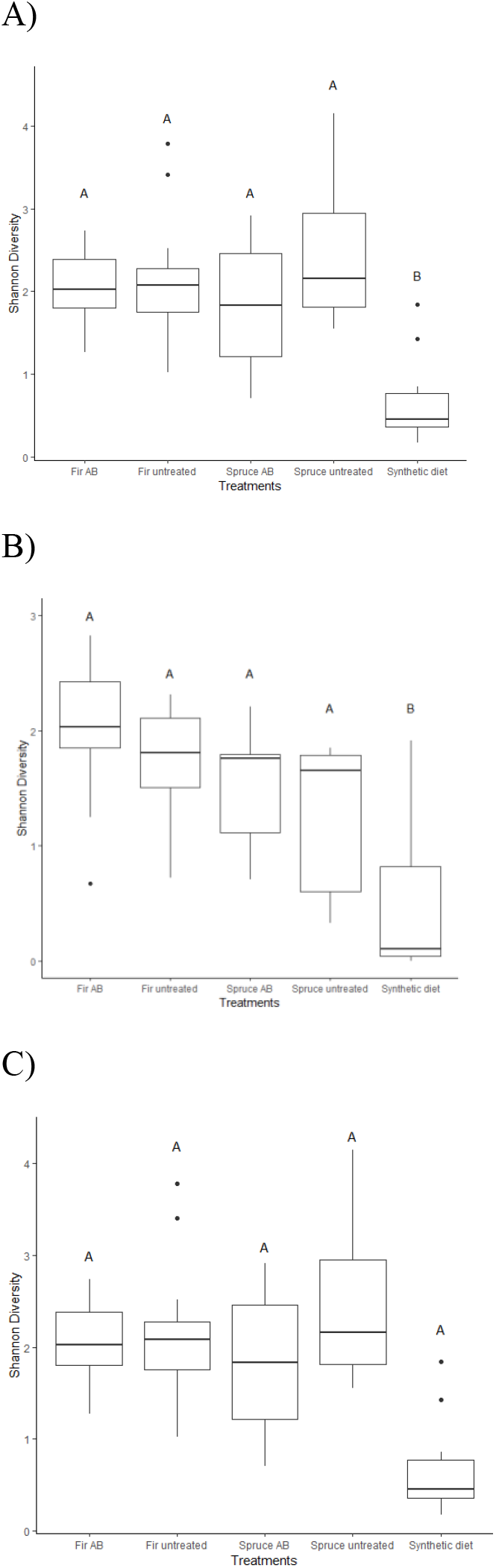
Mean (± SE) Shannon diversity of (A) spruce budworm diets (B) guts and (C) frass associated microbial communities among different diets (spruce versus fir foliage and synthetic diet) and antibiotic treatments (AB= antibiotic treated).

The effects of the antibiotic treatment were also observed between foliage samples (fir and spruce) when artificial diet was removed from the comparison. Foliage community structure was significantly altered by antibiotic treatment (PERMANOVA on weighted UniFrac; F=2.33, R^2^=0.06, p=0.024) but, there was no difference in the composition of foliage communities between antibiotic treatments (PERMANOVA on unweighted UniFrac; F=1.37, R^2^=0.38, p=0.096). We also observed a significant interaction between the effect of the type of diet and antibiotic treatment on the structure of the diet associated communities. This interaction was significant when comparisons were made among all diet samples (PERMANOVA on weighted UniFrac; F=3.05, R^2^=0.038, p=0.018) and between foliage samples (PERMANOVA on weighted UniFrac; F=2.35. R^2^=0.061, p=0.023).

We observed differences in diet associated community structure with antibiotic treatment, however we did not observe an effect of antibiotic treatment on the structure of spruce budworm gut microbial communities (PERMANOVA on weighted UniFrac; F=1.37, R^2^=0.019, p=0.21) (Fig. 3). In fact, antibiotic treatment did not affect the diversity or the structure of the microbial community in either spruce budworm guts or frass.

**Figure 3.**
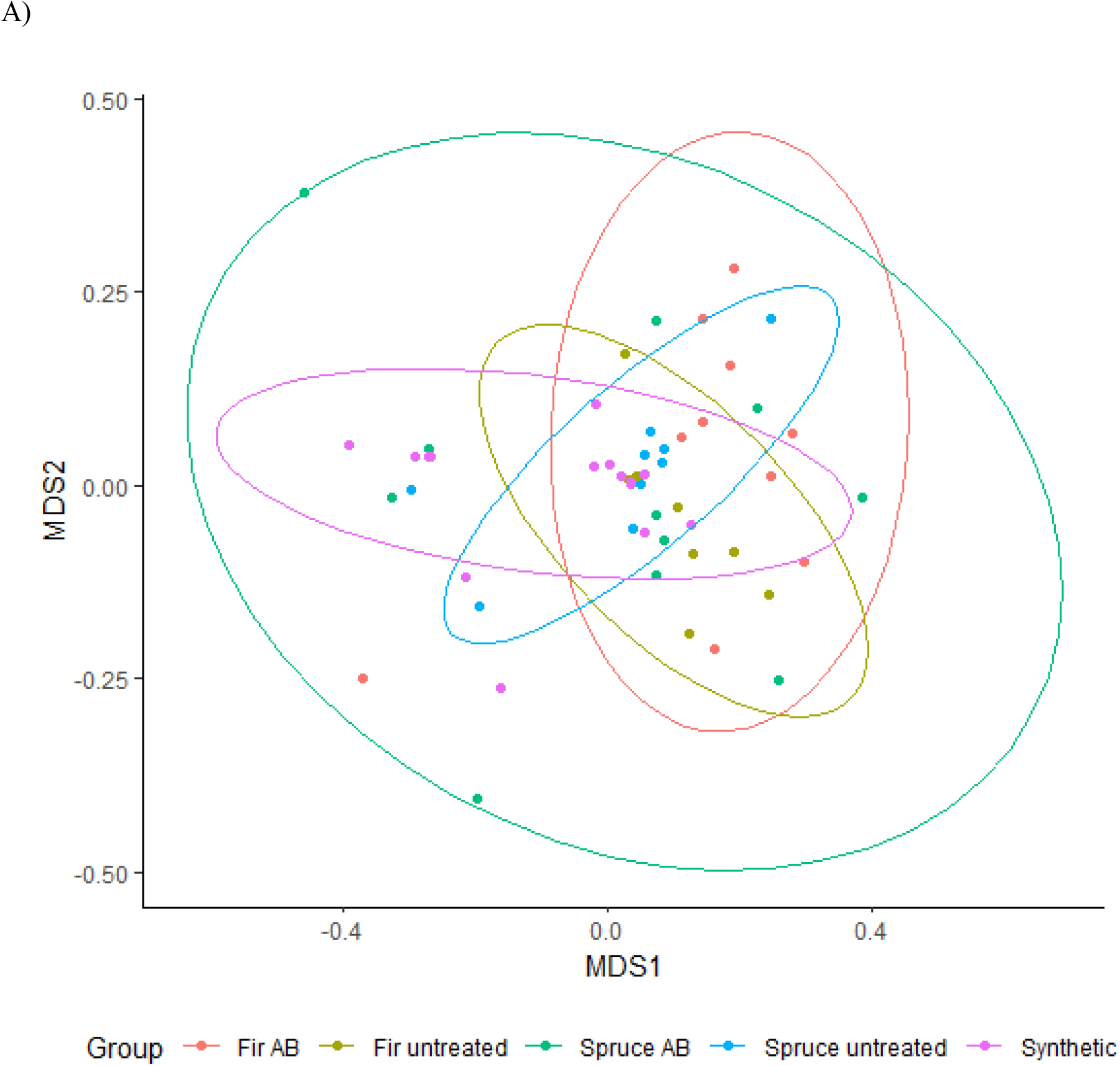
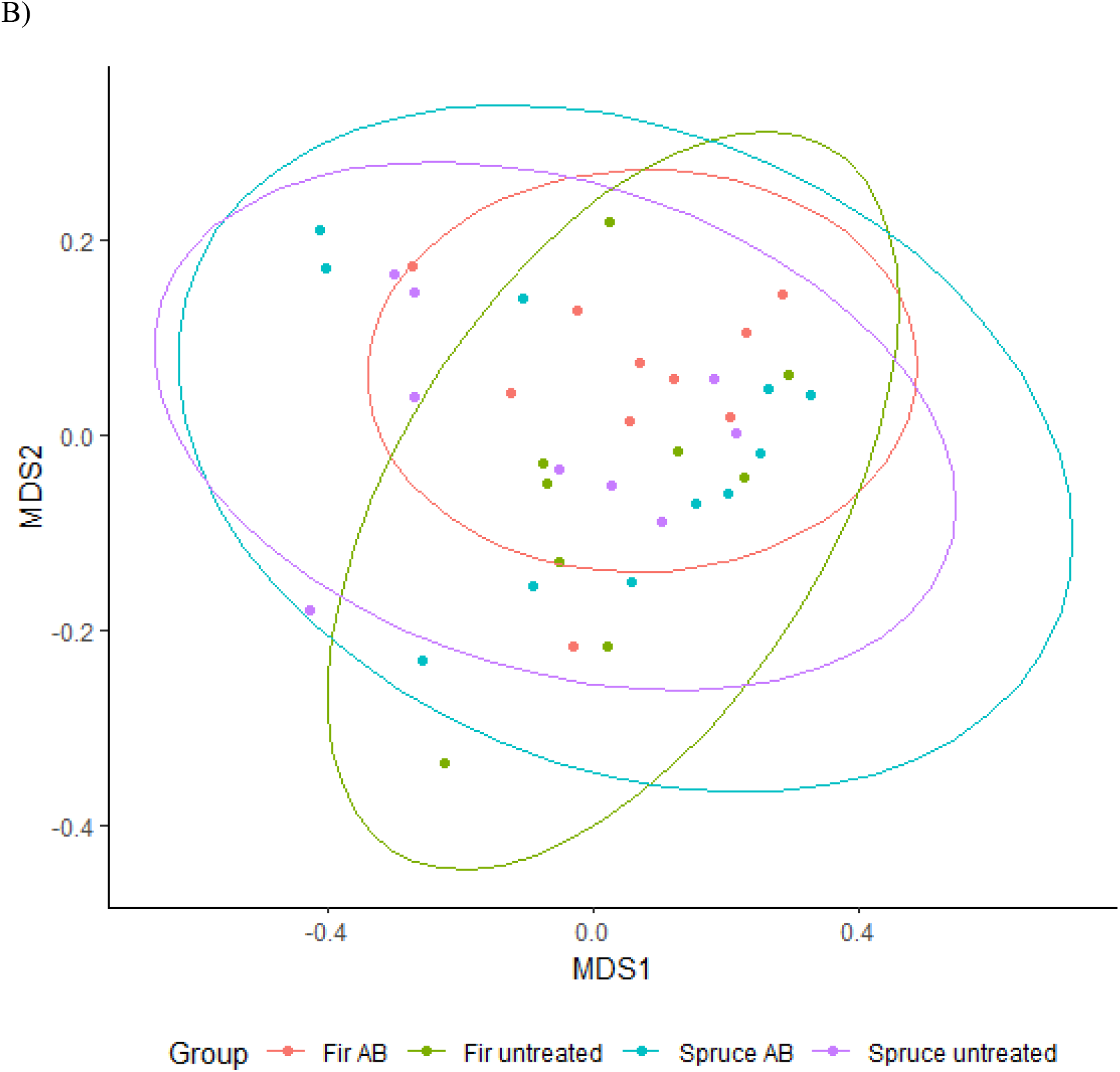
NMDS ordinations of gut-associated microbial communities based on weighted UniFrac distances. (A: all guts weighted UniFrac stress =0.07, B: guts of larvae feeding on foliage weighted UniFrac stress =0.07, Ellipses represent 95% confidence intervals around samples from different treatments (Fir.AB = antibiotic treated fir, Spruce.AB = antibiotic spruce, Fir.untreated = untreated fir, and Spruce.untreated = untreated spruce).

### Influence of dietary shifts on the microbial community independent of antibiotic treatment

In addition to antibiotic treatment, we included the diet type (i.e spruce foliage, fir foliage, or artificial diet) in our analysis. Here we explore the influence of host-tree species, or artificial laboratory diet, on the microbial community independent of antibiotics. The microbial diversity (Shannon diversity) was significantly different among diet types (ANOVA; F=20.47, p< 0.001)(Fig. 2). Synthetic diet, containing antibiotics, had lower Shannon diversity (± S.E) (0.65 ± 0.15) than either spruce (2.13 ± 0.22) or fir (2.13 ± 0.15) samples. Again, this difference in microbial diversity seems to be driven by the lower diversity in the synthetic diet because there was no significant change in microbial diversity between spruce and fir foliage samples (ANOVA; F= 0.00, p=0.993).

The type of diet also had a significant effect on the structure of diet-associated communities (PERMANOVA on weighted UniFrac; F=14.93, R^2^=0.378, p<0.001). This difference, however, was due to the differences between the synthetic diet and foliage. There was no difference in structure (PERMANOVA on weighted UniFrac; F= 1.89, R^2^=0.037, p=0.11) of foliage communities when synthetic diet communities were removed from the comparisons.

Guts of larvae that fed on synthetic diet had lower Shannon diversity (0.42 ± 0.15) on average than either the spruce fed (1.41 ± 0.12) or fir fed (1.85 ± 0.14) larvae (ANOVA; F=27.58, p<0.001)(Fig. 2). Unlike in diet samples we found that among foliage fed larvae, spruce fed larvae had lower bacterial diversity in their guts than larvae raised on fir (ANOVA; F=5.53, p=0.024). Diet had a significant effect on the gut microbial structure (PERMAOVA; F=8.95, R^2^=0.25, p<0.001) of spruce budworm larvae across all diets (Fig. 3).

The difference in gut community structure appears to be driven by the difference between synthetic diet-associated communities and foliage-associated communities because when the guts of only the larvae that were fed foliage were compared, there was no effect of foliage type (spruce vs fir) on gut community structure (Fig. 3–4). Finally, there was no effect of diet choice on the microbial diversity, community composition, or community structure of spruce budworm frass.

**Figure 4.**
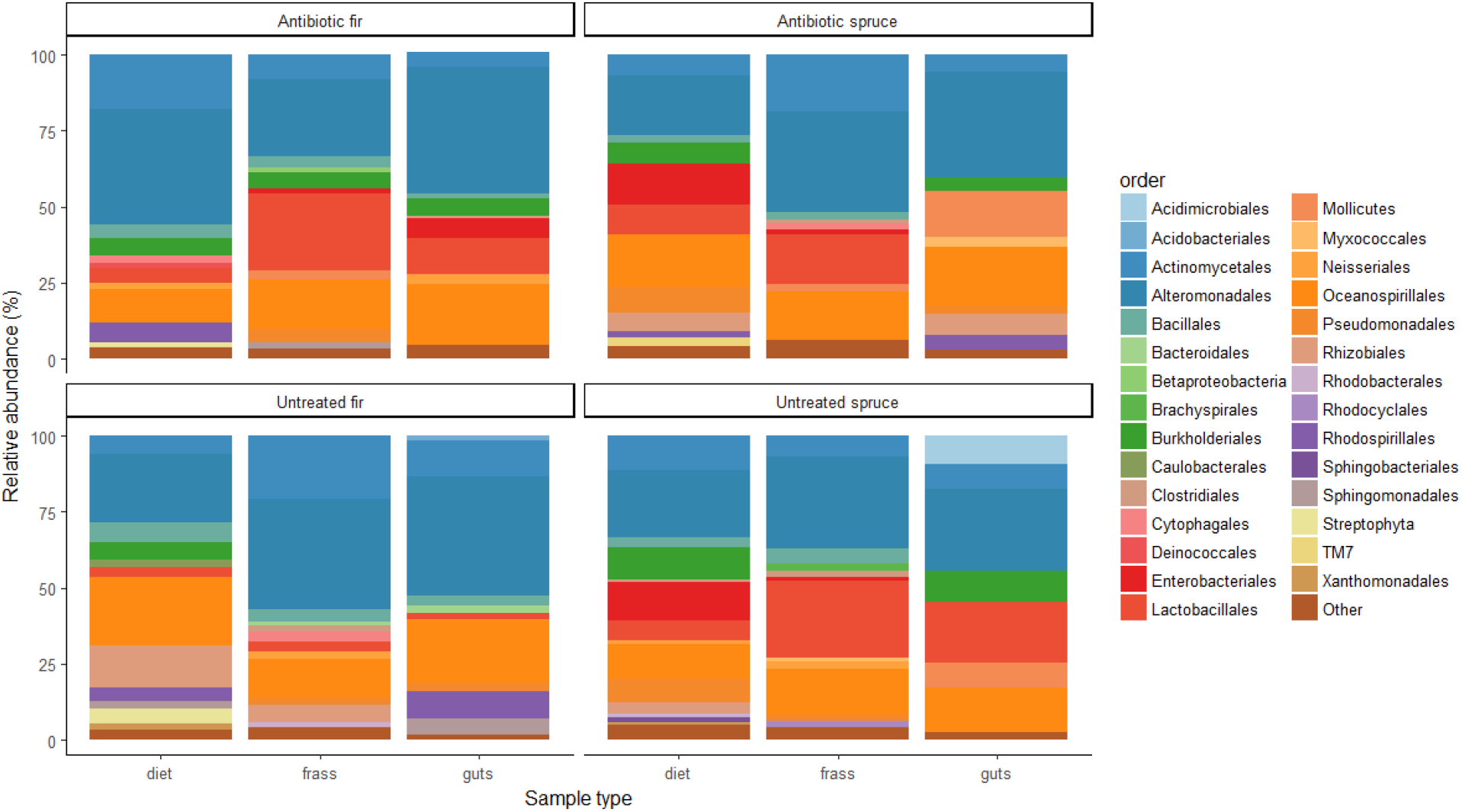
Mean relative abundance (%) of bacterial taxa across sample types, i.e diet (either spruce or fir foliage), budworm midguts, and budworm frass in each treatment group. Taxa were identified to the level of order, however if taxa remained unassigned at the order level they were labelled by taxonomic phylum or class.

Generally, our results show that the type of diet (spruce foliage, fir foliage, or synthetic diet) and antibiotic treatment had a significant effect on the communities associated with the diets fed to larvae from each treatment. When we analyzed the gut-associated bacterial communities we were still able to detect effects on community structure attributed to diet type but that difference was driven largely by the difference in community structure between synthetic diet and fresh foliage.

## DISCUSSION

We hypothesized that the use of antibiotics would reduce bacterial diversity as well as alter community composition and structure in the gut of the eastern spruce budworm. Contrary to our expectations, antibiotic treatment applied to the foliage that was given to larvae did not affect the bacterial diversity associated with the foliage-or gut-associated communities. Antibiotic treatment did, however, affect the structure of the foliage communities. This suggests that the antibiotic treatment did not affect the number of bacterial species present in the community but did affect the relative abundances of those taxa.

Our antibiotic treatment was sufficient to change the relative abundances of taxa present in foliage communities, but not to change the communities in the guts of spruce budworm larvae. Even if the effectiveness of the antibiotic treatment diminished as it was consumed alongside the food source, one would expect there to be some change in the gut-associated community structure compared to the control because the regional species pool (diet-associated community) is different between the antibiotic treated diets and the control diets. This suggests that there is either habitat filtering associated with the gut, likely the high alkalinity of the lepidopteran gut, or that the high alkalinity of the gut environment reduced the effectiveness of the antibiotic treatment. It is also possible that our dosage of antibiotics was not sufficient to influence the gut community. Because the measures of community diversity and structure used in this study were based on relative abundances, it is also possible that the antibiotics used in this study affected all bacteria equally thus reducing the total cell count but maintaining relative abundances. However, this remains to be evaluated using a measure of microbial abundance such as qPCR or flow cytometry.

### Influence of the gut microbial community on larval health

We further hypothesized that antibiotic treatment would result in a reduction of spruce budworm larval growth or survival. Despite not detecting structural or compositional shifts in the gut microbiome with antibiotic treatment our data suggest that the gut microbiome is not critical for growth. Considering that we found no differences in the communities of fast and slow growing larvae from the same treatment group, our results suggest that there is not a distinct microbial community associated with maintaining spruce budworm growth. We are confident that the spruce budworm gut microbiome is not critical in regulating growth. It is more likely that genetics and the nutrient quality of their food source are the principal drivers of spruce budworm larval growth.

Taken together with similar results from a growth experiment where the microbiome of *Manduca sexta* was eliminated via antibiotic treatment and no change in growth was detected (Hammer et al., 2017), our findings suggest that the eastern spruce budworm, and perhaps other lepidopteran species, are not nutritionally dependent on a microbial symbiosis. One possible explanation for this could be the bulk feeding strategy utilized by spruce budworm and many other herbivorous lepidopteran species. It is possible that because spruce budworm larvae consume so much food during their development, coupled with an extremely short gut retention time, it is not as imperative to efficiently extract nutrients from their diet. Another possible explanation could be that the alkalinity of the spruce budworm gut aids them in extracting nutrients or in tolerating the secondary compounds associated with conifer foliage without microbial assistance.

Along with previous work showing that carnivorous and herbivorous larval microbiomes did not differ significantly in composition (Whitaker, Salzman, Sanders, Kaltenpoth, & Pierce, 2016), this suggests that lepidopteran larvae may not select for gut bacteria based on their nutritional needs, providing further evidence lepidopteran larvae do not rely on microbial symbiosis to extract the necessary nutrients from food. An alternative interpretation of our results is that it is possible that the presence of a microbial community regulates growth or survival, but the composition or the relative abundance of its members is not important as seen in mosquitoes (Coon, Vogel, Brown, & Strand, 2014). It is also possible that the microbes in the gut are consumed for nutrients or as food, but they are either metabolically inactive or any metabolites produced are of no use to the host.

It should be noted that this experiment was on larvae with an already perturbed microbial community, since all larvae were fed the same synthetic diet containing antibiotics at the start of the experiment. This choice was to ensure a consistent starting point so that initial differences in larval microbiomes would not influence our results. We maintain that this does not invalidate our conclusion that the budworm gut microbial community does not influence host health for two reasons; first, although all larvae were fed with the synthetic diet, we maintained a control group on synthetic diet to ensure that our experimental treatments would change the starting community (which is characterized by our synthetic diet control group). Second, in our analysis we observed that communities associated with the artificial diet, or larvae reared on artificial diet, differed from foliage-associated communities. We feel that for these reasons our experimental design choice to rear all of our starting larvae on synthetic diet containing antibiotics does not invalidate our results. Either an experiment where field collected insects had their communities perturbed or an experiment using axenic insects could be used to validate these results.

Although we did not directly test whether the spruce budworm microbiome is resident or transient, our results shed some light on an ongoing discussion in the literature about the nature of gut microbiota in lepidopteran larvae (Hammer et al., 2017; Whitaker et al., 2016). Despite increasing evidence for habitat filtration by the host, we found that microbiome disturbance via antibiotics did not impact larval growth. These findings provide further evidence that lepidopteran larvae are not nutritionally dependent on microbial associations. It is possible, however, that other aspects of spruce budworm health could be influenced by the gut microbiome. It is also possible that the presence of bacteria in the gut are important to SBW larvae as an additional food source or for proper development of the innate immune system. More studies will be necessary to determine the influence of the microbiome on spruce budworm reproductive fitness, fecundity, and parasitism rates for example through field studies with wild populations of spruce budworm. It could also be possible for the gut microbiome to be more important in adult moths, however little is known about the microbiome in adult *C. fumiferana.*

### Conclusions

Overall, we observed that spruce budworm larvae do not appear to be nutritionally dependent on gut-associated microbiota for growth. We also found that spruce budworm larvae tended to have higher growth rates when feeding on a secondary host, black spruce, compared to balsam fir foliage. Our findings did not support our hypothesis that alteration of the spruce budworm gut-associated microbial community would reduce larval growth. We did however provide further evidence in support of the hypothesis that the gut microbial community of lepidopteran larvae is unimportant for larval growth.

## Acknowledgments

This work was supported by the FRQNT (Projet de recherche en équipe to S. Kembel, D. Kneeshaw, and P. James), NSERC (Discovery Grant Program), the Canada Research Chairs program, and SERG-I #: 2015/09-2018-908 (Project title: Using landscape level forest management and biotic interaction to reduce the intensity of spruce budworm outbreaks presented to S. Kembel, D. Kneeshaw, L. DeGrandpré, L. Kenefic, P. James, and D. Pureswaran)

We thank Mathieu Neau and Mathieu Landry for their help in sample collection, Tonia De Bellis for help with the production of figures, and Julie Marleau for technical assistance with the operation of the Illumina MiSeq.

