## Supplemental Figure 1 and Supplemental tables 1 and 2 for "The Gut Microbiome of the Eastern Spruce Budworm Does Not Influence Larval Growth or Survival"

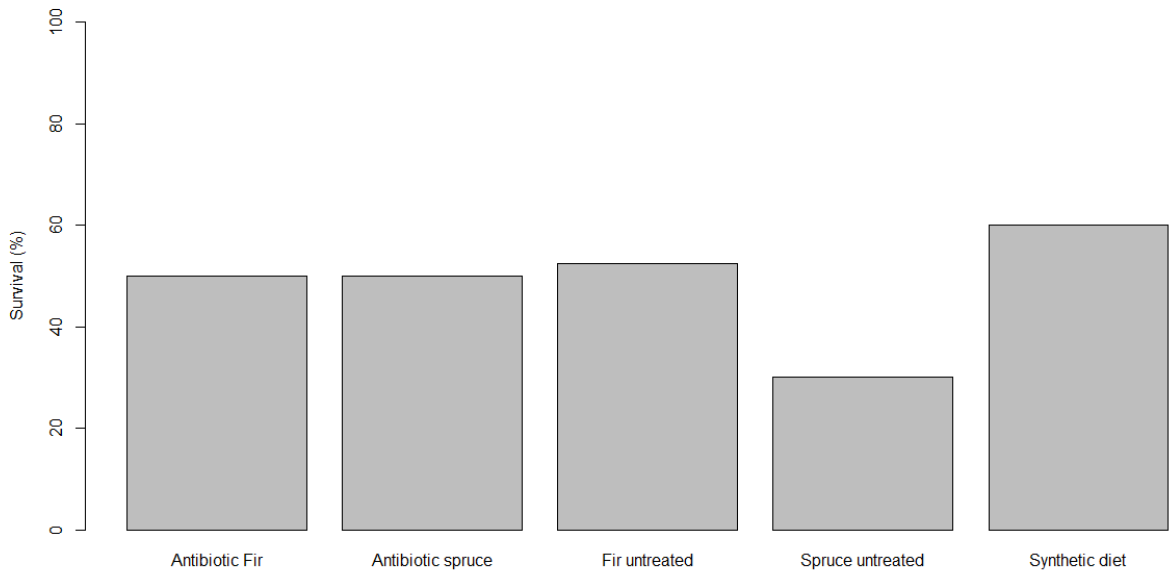


Figure S1. Percent survival of spruce budworm larvae among treatment diets (spruce versus fir foliage and synthetic diet) and antibiotic treatments (AB = antibiotic treatment).

Table S1. Estimates of the influence time, antibiotic treatment, and diet on the weight of spruce budworm larvae exposed to different diet and antibiotic treatments. Results represent ANOVA analysis of a mixed effect model with time, antibiotic treatment, and diet and their interactions as fixed factors and time nested within individual as random factors. In our model time as a fixed effect represents the growth rate of larvae and significant interactions between Time and other factors indicate that growth rates differed among treatment combinations.

| Variable | F value | P value |
|---|---|---|
| Intercept | 16637.5 | <.0001 |
| Time | 751.8 | <.0001 |
| Treatment | 5.78 | 0.018 |
| Diet | 3.2 | 0.076 |
| Time:Treatment | 3.0 | 0.081 |
| Time:Diet | 33.9 | <.0001 |
| Treatment:Diet | 3.8 | 0.056 |
| Time:Treatment:Diet | 2.1 | 0.151 |

Table S2. Pairwise comparisons of growth rate estimates for spruce budworm larvae within treatment groups as determined by a Tukey's Honest Significant Difference post-hoc test based on a mixed effect model with time, antibiotic treatment, and diet as fixed factors and time nested within individual as random factors.

| Contrast | Estimate | Standard error | Degrees of freedom | T ratio | P value |
|---|---|---|---|---|---|
| Antibiotic fir versus antibiotic spruce | -0.0290 | 0.0054 | 483 | -5.339 | <.0001 |
| Antibiotic fir versus untreated fir | -0.0012 | 0.0054 | 483 | -0.232 | 0.9956 |
| Antibiotic fir versus untreated spruce | -0.0184 | 0.0063 | 483 | -2.92 | 0.0191 |
| Antibiotic spruce versus untreated Fir | 0.0278 | 0.0053 | 483 | 5.165 | <.0001 |
| Antibiotic spruce versus untreated spruce | 0.0106 | 0.0062 | 483 | 1.693 | 0.3284 |
| Untreated fir versus untreated spruce | -0.0171 | 0.0062 | 483 | -2.744 | 0.0319 |
